# The effect of whey protein on viral infection and replication of SARS-CoV-2 and pangolin coronavirus in vitro

**DOI:** 10.1101/2020.08.17.254979

**Authors:** Huahao Fan, Yuqian Luo, Bixia Hong, Liqin Wang, Xiangshu Jin, Yangzhen Chen, Yunjia Hu, Tong Li, Hui Zhuang, Yi-Hua Zhou, Yigang Tong, Kuanhui Xiang

## Abstract

Since the detection of severe acute respiratory syndrome coronavirus 2 (SARS-CoV-2) in human breastmilk, little is known about the antiviral property of human breastmilk to SARS-CoV-2 and its related pangolin coronavirus (GX_P2V). Here we present for the first time that whey protein from human breastmilk effectively inhibited both SARS-CoV-2 and GX_P2V by blocking viral attachment, entry and even post-entry viral replication. Moreover, human whey protein inhibited infectious virus production proved by the plaque assay. We found that whey protein from different species such as cow and goat also showed anti-coronavirus properties. And commercial bovine milk also showed similar activity. Interestingly, the main antimicrobial components of breastmilk, such as Lactoferrin and IgA antibody, showed limited anti-coronavirus activity, indicating that other factors of breastmilk may play the important anti-coronavirus role. Taken together, we reported that whey protein inhibits SARS-CoV-2 and its related virus of GX_P2V. These results rule out whey protein as a direct-acting inhibitor of SARS-CoV-2 and GX_P2V infection and replication and further investigation of its molecular mechanism of action in the context of COVID-19.

## Introduction

Since the emergence of coronavirus disease 2019 (COVID-19) caused by severe acute respiratory syndrome coronavirus 2 (SARS-CoV-2) in late 2019, it imposes a great threat to global public health. COVID-19 resulted in high rate of infected patients and the collapse of health systems in several countries(1, 2). It is reported by world Health Organization (WHO) that more than 18 million people are SARS-CoV-2 positive and over 700 thousand people died by COVID-19.

As a novel pathogen in human, SARS-CoV-2 could transmit and lead to disease among various population, including pregnant women(1, 3, 4). Despite no evidence to support the vertical transmission of SARS-CoV-2, it has been reported that SARS-CoV-2 RNA could be detected in human breastmilk(3, 5). Considering that breastmilk can inhibit some viruses, such as human immunodeficiency virus (HIV), cytomegalovirus (CMV) and dengue virus(6, 7), we supposed that breastmilk could also inhibit SARS-CoV-2 infection.

It is well known that milk has a rich source of biologically active components of valuable proteins, minerals and vitamins. Interestingly, the protein fraction has many kinds of biological functions. In particular, milk has antibacterial and antiviral properties(7). It was reported that milk components show antiviral activity against the HIV, HCMV and hepatitis C virus (HCV)(8–10). The Lactoferrin from the milk was reported to show antiviral activity and inhibition of intracellular HCV replication(11, 12). However, it is still unknown whether human breastmilk show anti-SARS-CoV-2 activity and protect the newborn from SARS-CoV-2 infection.

Here, we performed the study of the inhibition of SARS-CoV-2 and its related pangolin coronavirus (GX_P2V) by whey protein in cultured cell lines.

## Material and methods

### Collecting and handling of milk samples

Breastmilk was collected via pumps into sterile containers after disinfecting nipples with 75% ethanol. Samples were frozen in aliquots at −80°C. Skim milk was prepared by centrifuging the samples for 15 min at 4,000 × g at 4°C and the lower aqueous phase was used for further experiments and analysis. Mothers were informed consent. They were not infected with hepatitis B and C viruses and HIV. This study was approved by the ethics committees of the Medical Center.

### Cell lines, coronavirus, and key reagents

Vero E6 cells (American Type Culture Collection, Manassas, VA, USA) and A549 cells were grown in high-glucose-containing Dulbecco's Modified Eagle Medium supplemented with 10% fetal bovine serum(13). SARS-CoV-2 pseudovirus was kindly shared by Prof. Youchun Wang (National Institutes for Food and Drug Control, China)(14). GX_P2V was described recently(15). Goat and cow whey protein were purchased from Sigma (USA). The recombined lactoferrin (rLF), human lactoferrin (hLF) and bovine lactoferrin (bLF) were purchased from Sigma (USA).

### Viral infection assay

SARS-CoV-2 pseudovirus infection was performed as described(14). The cells were infected with viral inocula of 650 TCID50/well. One day post infection (1dpi), the cells were lysed and the luminescence was measured according to the manufacture’s protocol. Cells were infected with GX_P2V at multiplicity of infection (MOI) of 0.01 as described(13). The messenger RNA (mRNA) levels of GX_P2V and GAPDH were determined by reverse transcription and quantitative real-time polymerase chain reaction (RT-qPCR)(13).

### Plaque assay for determining virus titer

The plaque assay for determining virus titer was performed as previously described(13). Briefly, confluent monolayer Vero E6 cells were infected with GX_P2V with ten-fold dilution from 10^−1^ to 10^−6^. After removing the virus, the cells were washed by PBS and added with 1% agarose overlay to prevent cross contamination. At 5 dpi, cells were fixed with 4% paraformaldehyde for 1h, followed by staining with Crystal violet for 10 min and washed with water. The plaques were counted and virus titers were calculated.

### Blocking assay

To study whether the inhibition effects of skimmed breastmilk on SARS-CoV-2 pseudovirus and GX_P2V by binding cell surface to block viral entry, Vero E6 cells were seeded at the 96 well plate with 20,000 cells/well. The skimmed breastmilk at a final concentration of 4mg/ml was added in the wells. After washing away the free breastmilk the next day, the cells were infected with SARS-CoV-2 pseudovirus and cultured for 24h. The luminescence was measured to reflect the viral infection and replication.

### Viral attachment assay

Vero E6 cells were seeded at the 96 well plate with 20,000 cells/well one day before the infection. Breastmilk at a final concentration of 4mg/ml was mixed with SARS-CoV-2 pseudovirus (650 TCID_50_/well) and GX_P2V (MOI=10) at 4°C for 1h. The mixture was added into the cells and put at 4°C for 2h to allow viral attachment to cells. After washing out of free virus, cell surface GX_P2V was extracted and quantified by RT-qPCR(16). For pseudovirus, the free viruses were washed away and incubated at 37°C for 24h. The luminescence was measured to reflect the viral infection and replication.

### Viral entry assay

Breastmilk at a final concentration of 4mg/ml was mixed with SARS-CoV-2 pseudovirus (650 TCID_50_/well) and GX_P2V (MOI=10) at 4 °C for 1h. Vero E6 cells were exposed to the mixture at 37°C for 1h to allow viral internalization into cells. GX_P2V mRNA was measured by RT-qPCR. For pseudovirus, the free viruses were washed away and the cells were incubated at 37°C for 24h. The luminescence was measured.

### Viral post-entry assay

Vero E6 cells were infected with SARS-CoV-2 pseudovirus (650 TCID_50_/well) and GX_P2V (MOI=0.01) and incubated at 37°C for 1h, respectively. After washing out of the free viruses, the cells were cultured in the media containing breastmilk at a final concentration of 4mg/ml for 24h (SARS-CoV-2 pseudovirus) and 72h (GX_P2V), respectively. Intracellular SARS-CoV-2 pseudovirus was measured by luminescence and GX_P2V was measured by RT-qPCR.

### Viral RNA extraction and quantification

The RT-qPCR for GX_P2V RNA quantification was performed as previously described(13). Briefly, total RNA was extracted by the AxyPrep™ multisource total RNA Miniprep kit (Axygene, USA). First strand complementary DNA (cDNA) was synthesized by a Hifair II 1st Strand cDNA synthesis kit with gDNA digester (Yeasen Biotech, China) and quantified by Hieff qPCR SYBR Green Master Mix (Yeasen Biotech, China). The primer sequences were listed in **Supplementary Table 1**. The RT-qPCR amplification of the Taqman method was performed as follows: 50 °C for 2 min, 95 °C for 10 min followed by 40 cycles consisting of 95 °C for 10 s, 60 °C for 1 min.

### Western blotting

Western blotting was performed as described previously(17). Briefly, the samples were loaded on a 12% SDS-PAGE gel and transferred to a polyvinylidene fluoride membrane. Antibody against nucleocapsid protein of anti-SARS-CoV-2 N protein (Genscript, USA) and GAPDH of anti-GAPDH (Proteintech, USA) were used at 1:3000 dilutions. The second antibody of HRP-conjugated affinipure Goat anti-mouse IgG (H+L) were diluted at 1:20000. SuperSignal^®^ West Femto Maximum Sensitivity Chemiluminescent Substrate (Thermo Scientific, USA) was used for imaging.

### Statistical analysis

Statistical analyses were analyzed using GraphPad Prism 8 software (GraphPad Software Inc., San Diego, CA, USA). Values are shown as mean of triplicates. Comparisons between the two groups were analyzed using the Student's t tests. Values of *p*<0.05 was considered statistically significant.

## Results

### Inhibition of SARS-CoV-2 pseudovirus by human breastmilk

To determine the potential inhibition of SARS-CoV-2 by human breastmilk, we infected the Vero E6 cells with SARS-CoV-2 pseudovirus of 650 TCID_50_/well in the presence of human skim breastmilk (4mg/ml) until the end of the experiment. At 24 hours post infection (hpi), we harvested the cells and monitored the luciferase expression, which reflects the infection of SARS-CoV-2. As shown in Fig.1A, all the skim breastmilk from different donors effectively inhibited SARS-CoV-2 pseudovirus with inhibition efficiency more than 98%.

To confirm the inhibition of SARS-CoV-2 by human breastmilk, we did serial 2-fold dilution of the breastmilk and tested their inhibition. As shown in Fig.1B-1H, the inhibition efficiency of breastmilk was doses dependent. Breastmilk could inhibit SARS-CoV-2 at a low concentration with different concentration for 50% of maximal effect (EC_50_) of the donors: N17A (EC_50_=0.25mg/ml) (Fig.1B), N30 (EC_50_=0.29mg/ml) (Fig.1C), N45 (EC_50_=0.02mg/ml) (Fig.1D), N49 (EC_50_=0.16mg/ml) (Fig.1E), N62 (EC_50_=0.02mg/ml) (Fig.1F), N66 (EC_50_=0.05mg/ml) (Fig.1G), N68 (EC_50_=0.25mg/ml) (Fig.1H), indicating that the concentration of the effective factors for SARS-CoV-2 inhibition showed individual difference. Interestingly, breastmilk showed low cytotoxicity with cytotoxicity concentration 50% (CC_50_) more than 3mg/ml. These results indicated that human breastmilk showed high anti-SARS-CoV-2 property.

**Fig.1.**
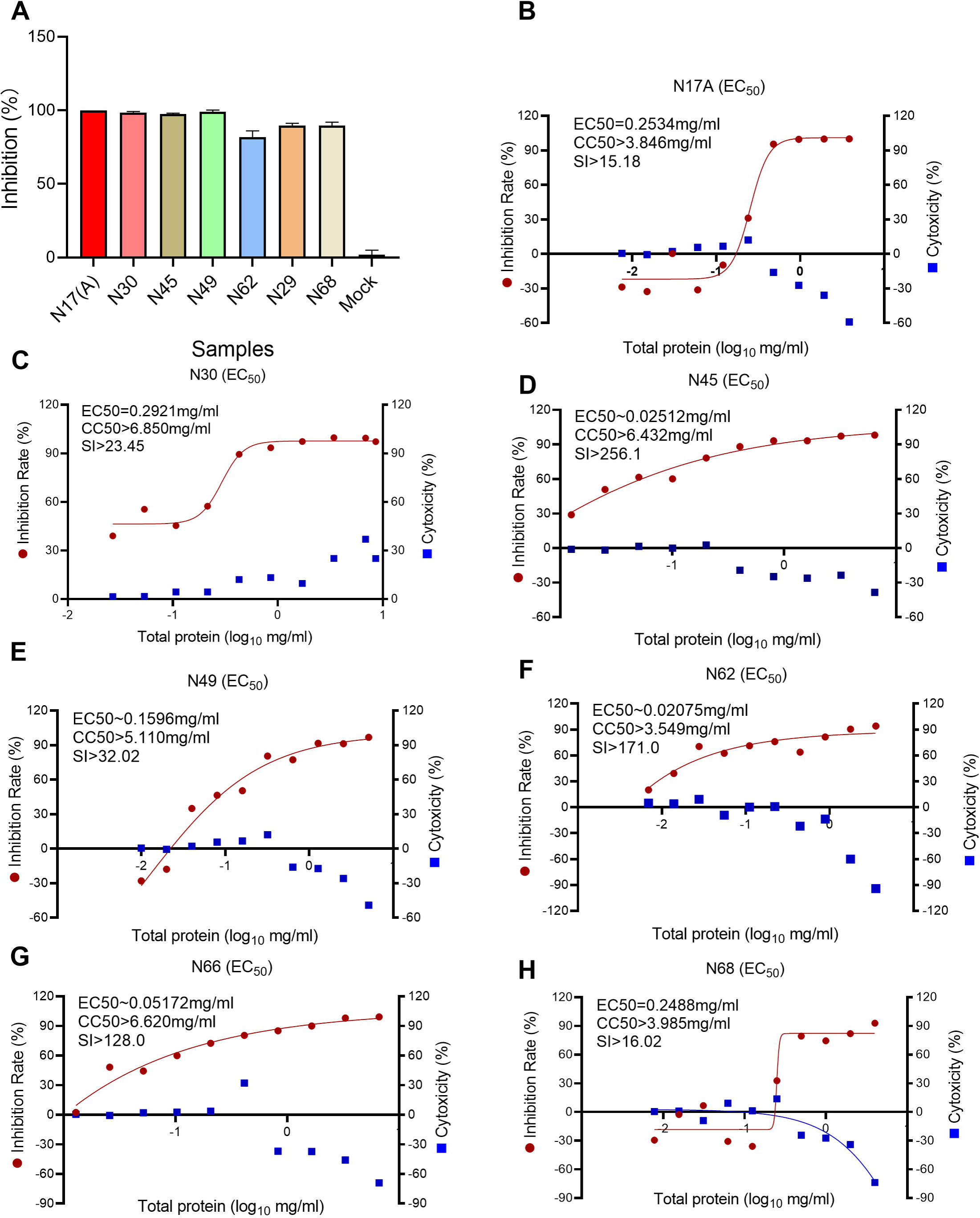
Inhibition of SARS-CoV-2 pseudovirus by human breastmilk. (A) Vero E6 cells were infected with SARS-CoV-2 pseudovirus (650 TCID_50_/well) in the present of breastmilk from different donors with final concentration of 4mg/ml for 24h. Luciferase reflecting SARS-CoV-2 infection were detected by the microplate luminometer. Different doses of breastmilk from donors of N17A (B), N30 (C), N45 (D), N49 (E), N62 (F), N66 (G) and N68 (H) were used to treat SARS-CoV-2 pseudovirus infection of Vero E6 cells for 24h. The luciferase in the cells was quantified by the microplate luminometer. Cytotoxicity of these drugs to Vero E6 cells was measured by CellTiter-Blue assay. The left and right Y-axis of the graphs represent mean percentage of inhibition of virus yield and cytotoxicity of breastmilk, respectively. Values are shown as mean of triplicates.

### Inhibition of GX_P2V by human breastmilk

As reported recently, the SARS-CoV-2 related virus of GX_P2V shares 92.2% amino acid identity in spike protein with SARS-CoV-2, which is a suitable model for SARS-CoV-2 infection research(13). In this study, we also utilized GX_P2V to study the inhibition of SARS-CoV-2 by human breastmilk. Vero E6 cells were infected with mixtue of GX_P2V virus (MOI=0.01) and human breastmilk (4mg/ml). As shown in Fig. 2A, all the breastmilk from different donors showed nearly 100% of inhibition on the infectivity of GX_P2V.

Next, we did serial 2-fold dilution of human breastmilk and tested their inhibition effects. Similar to the inhibition of SARS-CoV-2 pseudovirus, GX_P2V virus was inhibited doses dependently by human breastmilk. The EC_50_ of breastmilk from different donors were different: N30 (EC_50_=0.57mg/ml) (Fig.2B), N45 (EC_50_=0.34mg/ml) (Fig.2C), N62 (EC_50_=0.84mg/ml)(Fig.2), N66 (EC_50_=0.79mg/ml) (Fig.2E) and N68 (EC_50_=0.67mg/ml) (Fig.2F). Interestingly, human breastmilk didn’t show any cytotoxicity to Vero E6 cells (CC_50_>3mg/ml), and even promoted cells proliferation. In all, human breastmilk could also inhibit SARS-CoV-2 related virus of GX_P2V.

**Fig.2.**
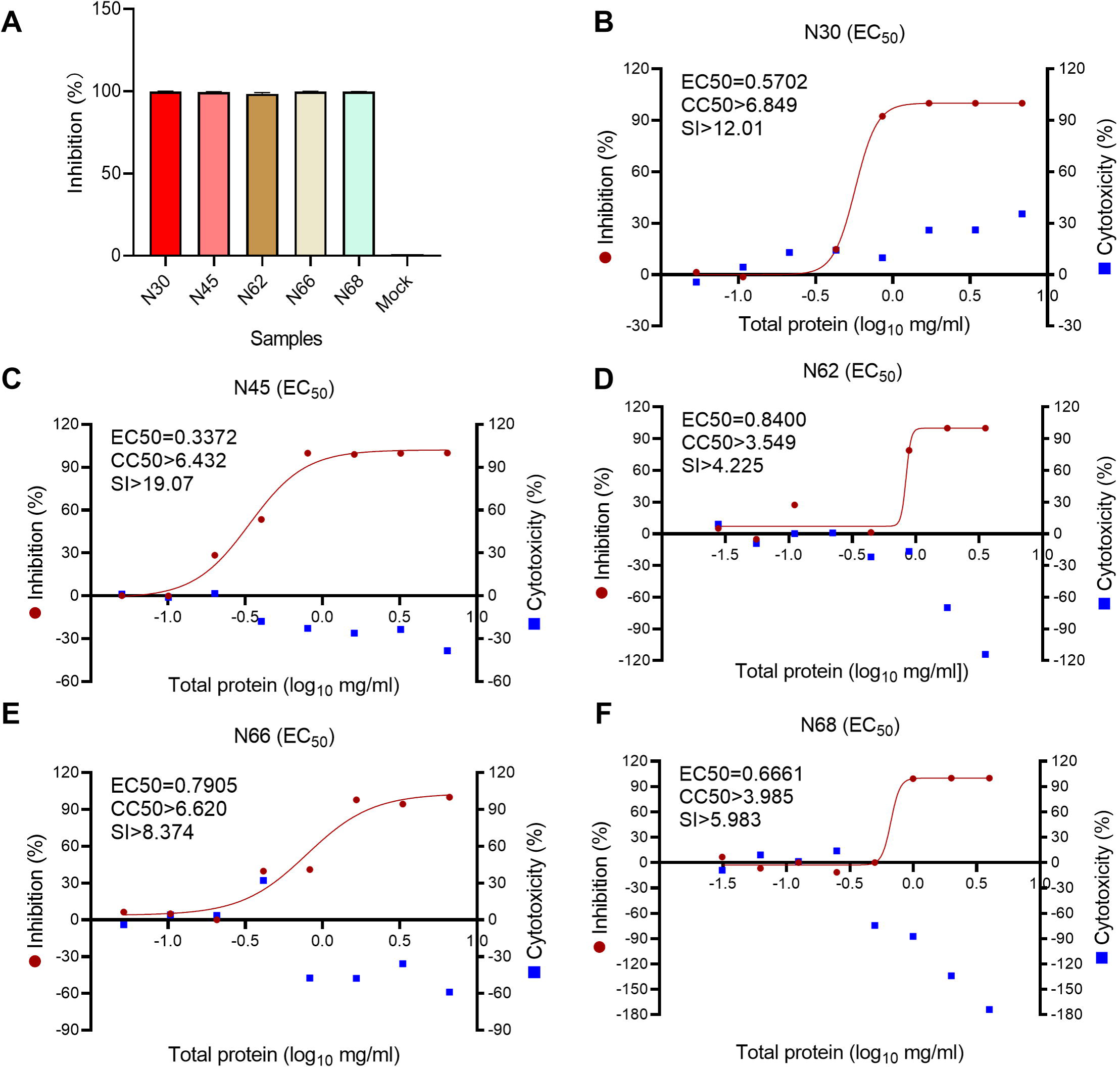
Inhibition of GX_P2V infection by human breastmilk. (A) Vero E6 cells were infected with GX_P2V (MOI=0.01) in the presence of breastmilk from different donors with final concentration of 4mg/ml for 72h. GX_P2V RNA level were measured by RT-qPCR and normalized by GAPDH level. Different doses of human breastmilk from donor of N30 (B), N45 (C), N62 (D), N66 (E) and N68 (F) were tested to inhibit GX_P2V. Cytotoxicity of these samples to Vero E6 cells was measured by CellTiter-Blue assay. The left and right Y-axis of the graphs represent mean percentage of inhibition of virus yield and cytotoxicity of the samples, respectively. Values are shown as mean of triplicates.

### Inhibition of infectious virus production by human breastmilk

To investigate the impact of human breastmilk on infectious virus production, experiments were performed. Vero E6 cells were infected by GX_P2V at a MOI of 0.01 in the presence of human breastmilk at concentration of 4, 0.8 and 0.16mg/ml, respectively. As shown in Fig.3A, after 72 hpi, GX_P2V RNA level increased during the dilution of breastmilk. Similarly, western blotting assay showed that GX_P2V nucleocapsid protein expression was highest in 0.16 mg/ml of breastmilk treatment but lowest in 4mg/ml of breastmilk (Fig.3B). These results were further proved by the inhibition of GX_P2V virus. Then, we did serial 10-fold dilution of the supernatant from the breastmilk treated group and infected the Vero E6 cells seeded in the dishes. After 5 dpi, we did the plaque assays. As shown in Fig.3C, the plaque assays showed that live viruses were positive in all the dishes and gradually decreased during the dilution of control group. However, when treated by 4mg/ml of breastmilk, all the dishes showed no live viruses. The live viruses could be found in the group of 0.16 mg/ml breastmilk treatment at the dilution of supernatant less than 1000 folds. The dilution of virus titers from 0.16 mg/ml breastmilk treatment less than 100000 folds still showed infectious virus. These results confirmed that breastmilk also inhibit SARS-CoV-2 related coronavirus infection.

**Fig.3.**
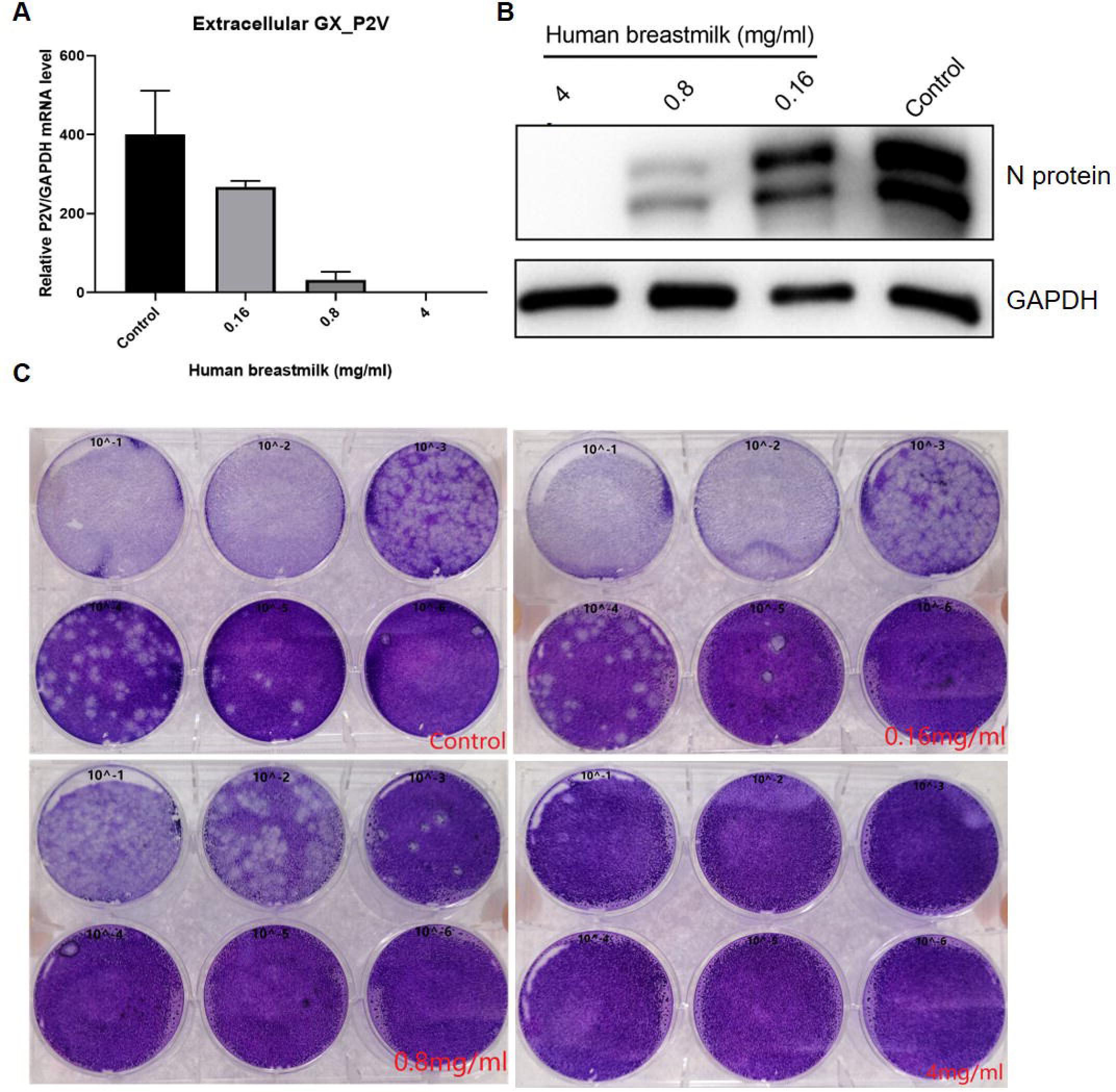
Human breastmilk inhibited infectious virus production. (A) Vero E6 cells were infected GX_P2V virus at a MOI of 0.01 in the treatment of different doses of human breastmilk for 72h. (A) GX_P2V RNA was quantified by RT-qPCR and normalized by GAPDH level. (B) GX_P2V NP was detected by western blot. (C) The virus titers of supernatant of control and different concentration of human breastmilk treatment were determined by plaque assay. MOI, multiplicity of infection; RT-qPCR, Real time quantitative polymerase chain reaction. Values are shown as mean of triplicates ± SD, \*\**p*<0.01, \*\*\**p*<0.001 by unpaired two-tailed t test.

### The inhibition of SARS-CoV-2 by whey protein from different species

To rule out whether whey protein in breastmilk play the important role in the inhibition of viral infection, we inactivated the protein by high temperature of 90 °C for 10 mins and protease K digestion, respectively, and test their role in viral inhibition. As shown in Fig.4A, the inhibition rate in breastmilk with treatment by heat and protease K showed limited inhibition of viral infection, indicating that whey protein none other factors in breastmilk has antiviral properties. Similar results were showed in A549 cell line (Fig.4B).

To investigate if whey protein from other species could also inhibit SARS-CoV-2 and GX-P2V infection, we selected cow and goat whey protein and performed the experiments. The Vero E6 cells were infected with SARS-CoV-2 pseudovirus at a MOI of 650 TCID_50_/well treated with serial 2-fold dilution of commercial bovine milk (Aptamil, Australia). As shown in Fig.4C, SARS-CoV-2 pseudovirus infection could be inhibited by commercial bovine milk doses dependently. When treated with human skimmed breastmilk, cow and goat whey protein at concentration of 4mg/ml, SARS-CoV-2 pseudovirus infection was also inhibited by whey protein from all the species. The inhibition by human skimmed breastmilk was significantly higher than that by cow and goat whey protein (Fig.4D). Similar results were observed in the GX_P2V infection experiments (Fig.4E). These results indicated that breastmilk from different species could inhibit SARS-CoV-2 infection, but with different inhibitory efficiency.

**Fig.4.**
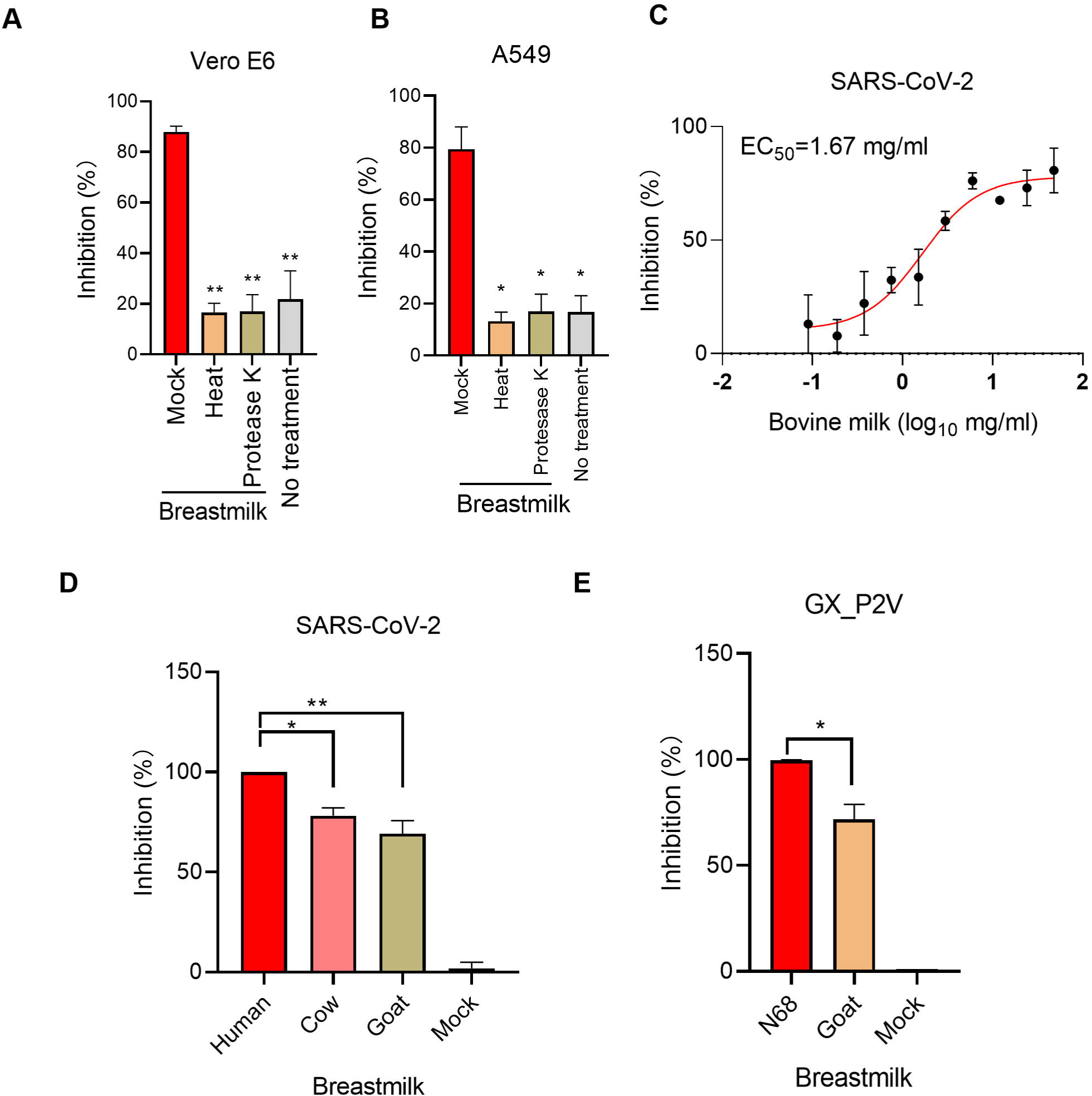
The inhibition of whey protein from different species were different. Whey protein not fatty showed the inhibition property. (A) Vero E6 cells and (B) A549 cells were infected with SARS-CoV-2 pseudovirus treated with human breastmilk with or without inactivation by heat or protease K. (C) The skimmed commercial bovine milk with different doses was used to treat SARS-CoV-2 pseudovirus for 24h. (D) Vero E6 cells were infected with SARS-CoV-2 pseudovirus with treatment by whey protein (4mg/ml) from human, cow and goat. Luciferase reflecting SARS-CoV-2 infection were detected by the microplate luminometer. (E) Vero E6 cells were infected with GX_P2V (MOI=0.01) in the present of human and goat whey protein at concentration of 4mg/ml for 72h. GX_P2V RNA level were measured by RT-qPCR and normalized by GAPDH level. Values are shown as mean of triplicates ± SD, \*\**p*<0.01, \*\*\**p*<0.001 by unpaired two-tailed t test.

### The impact of Lactoferrin and IgA antibody on SARS-CoV-2 virus

It is reported that lactoferrin has broad anti-viral effects(7). To determine if the inhibition of SARS-CoV-2 virus by lactoferrin, we treated the cells with virus and each of recombinant lactoferrin (rLF), bovine lactoferrin (bLF) and human lactoferrin (hLF) at concentration of 1mg/ml, respectively. Human breastmilk (1mg/ml) was used as control. As shown in Fig.5A, all the LFs showed limited inhibition to SARS-CoV-2 virus compared to the skimmed human breastmilk. Similarly, lactoferrin showed limited inhibition of GX_P2V (Fig.5B), indicated that there would be other factors in milk inhibiting SARS-CoV-2 virus.

It also reported that IgA antibody from recovered COVID-19 patients inhibited SARS-CoV-2 infection *in vitro*(18). However, the breastmilk we collected was in 2017, which mothers didn’t experience the COVID-19. To exclude the potential IgA antibody impact on SARS-CoV-2 infection, we utilized the neutralized anti-IgA antibody (1mg/ml) with different dilution to mix with the breastmilk (1mg/ml). As shown in Fig.5C, the different dilution of anti-IgA antibody didn’t influence the SARS-CoV-2 infection, indicating that IgA antibody from breastmilk is not the key factor inhibiting viral infection.

**Fig. 5.**
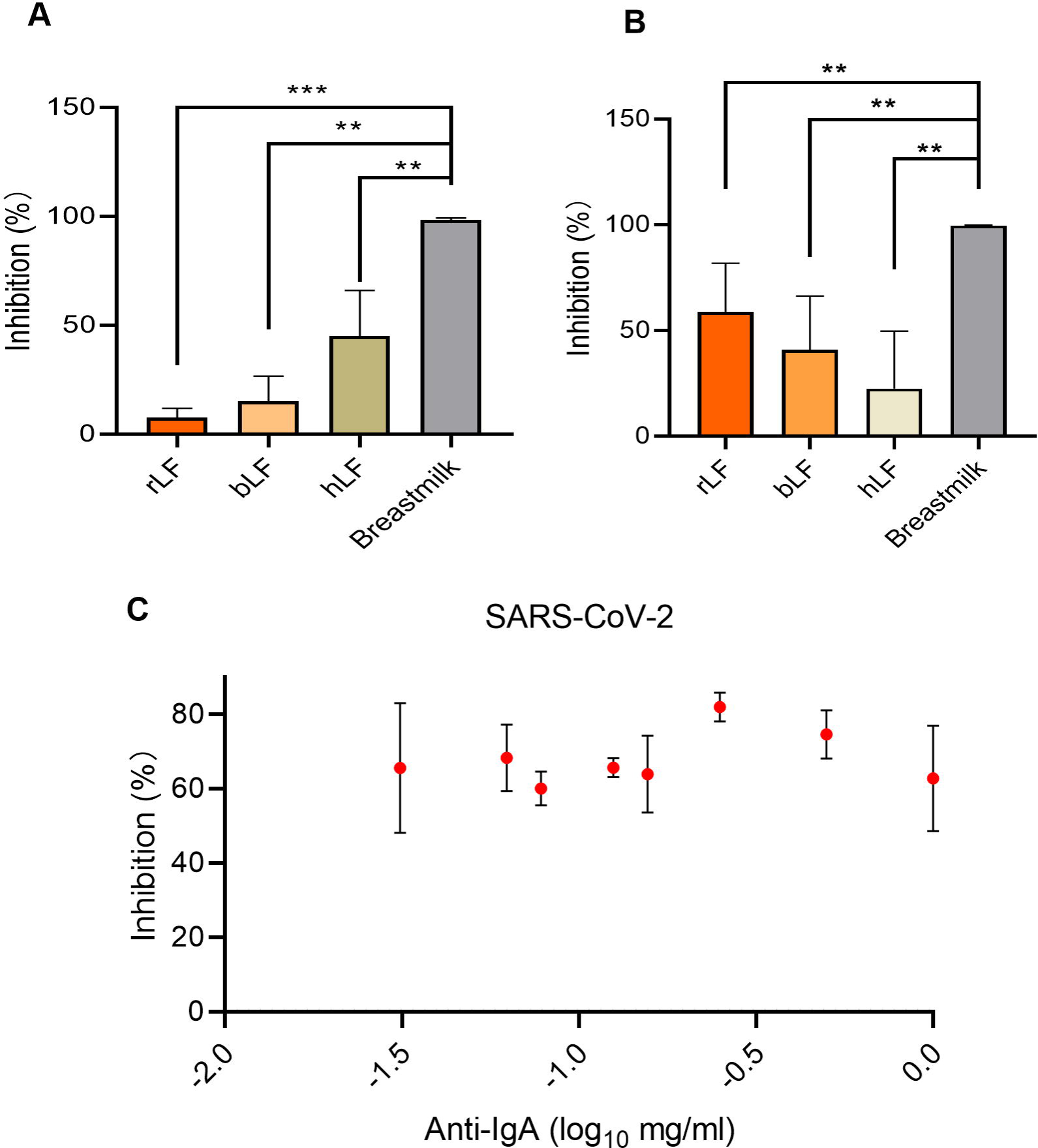
Lactoferrin and IgA antibody might not be the main factor for viral inhibition. Vero E6 cells were infected with (**A**) SARS-CoV-2 pseudovirus (650 TCID_50_/well) and (**B**) GX_P2V (MOI=0.01) in the presence of rLF, bLF and hLF at concentration of 1 mg/ml. Luciferase reflecting SARS-CoV-2 infection were detected by the microplate luminometer. (**C**) Anti-IgA antibody with different doses were used to neutralize the IgA and test their inhibition of IgA to SARS-CoV-2 pseudovirus. Luciferase was detected by the microplate luminometer. Values are shown as mean of triplicates ± SD, \*\**p*<0.01, \*\*\**p*<0.001 by unpaired two-tailed t test. rLF: recombined lactoferrin; bLF: bovine lactoferrin; hLF: human lactoferrin.

### The impact of whey protein on viral attachment, entry and post-entry replication

To further determine which step in the coronavirus life cycle was affected by whey protein of human breastmilk, we performed following experiments. Generally, SARS-CoV-2 attaches to cells and subsequently internalizes into cells through endocytosis, which leads to intracellular viral replication. The attachment of GX_P2V was significantly decreased in the whey protein treated cells (Fig.6A). These data were consistent with the results of SARS-CoV-2 pseudovirus (Fig.6E), two viruses share the similar mechanisms during the viral internalization process.

To examine the process after attachment, we mixed the breastmilk with GX_P2V for 1h at room temperature. Vero E6 cells were incubated with the mixture at 37 ºC for 1h. Then we harvested the cells and detected the viral RNA by RT-qPCR and normalized by GAPDH level. As shown in Fig.6B, the entry process of GX_P2V was significantly inhibited by breastmilk, which was consistent with the SARS-CoV-2 pseudovirus results (Fig.6F). To examine the post-entry replication process, we infected the cells with GX_P2V at 37 ºC for 1h, then removed the supernatant and treated the cells with breastmilk (4mg/ml). The GX_P2V level showed that human breastmilk could also inhibit post-entry viral replication (Fig.6C). In addition, to investigate the blocking activity of human whey protein on SARS-CoV-2, we mixed the breastmilk before the viral infection. Then, we washed the cells with PBS 5 times and infected SARS-CoV-2 pseudovirus for 24h. As shown in Fig.6D, both breastmilk treatment before and during the infection showed high inhibition efficiency to SARS-CoV-2. These results clearly suggested that human breastmilk plays a significant inhibition role in SARS-CoV-2 and related virus (GX_P2V) attachment, entry and replication.

**Fig.6.**
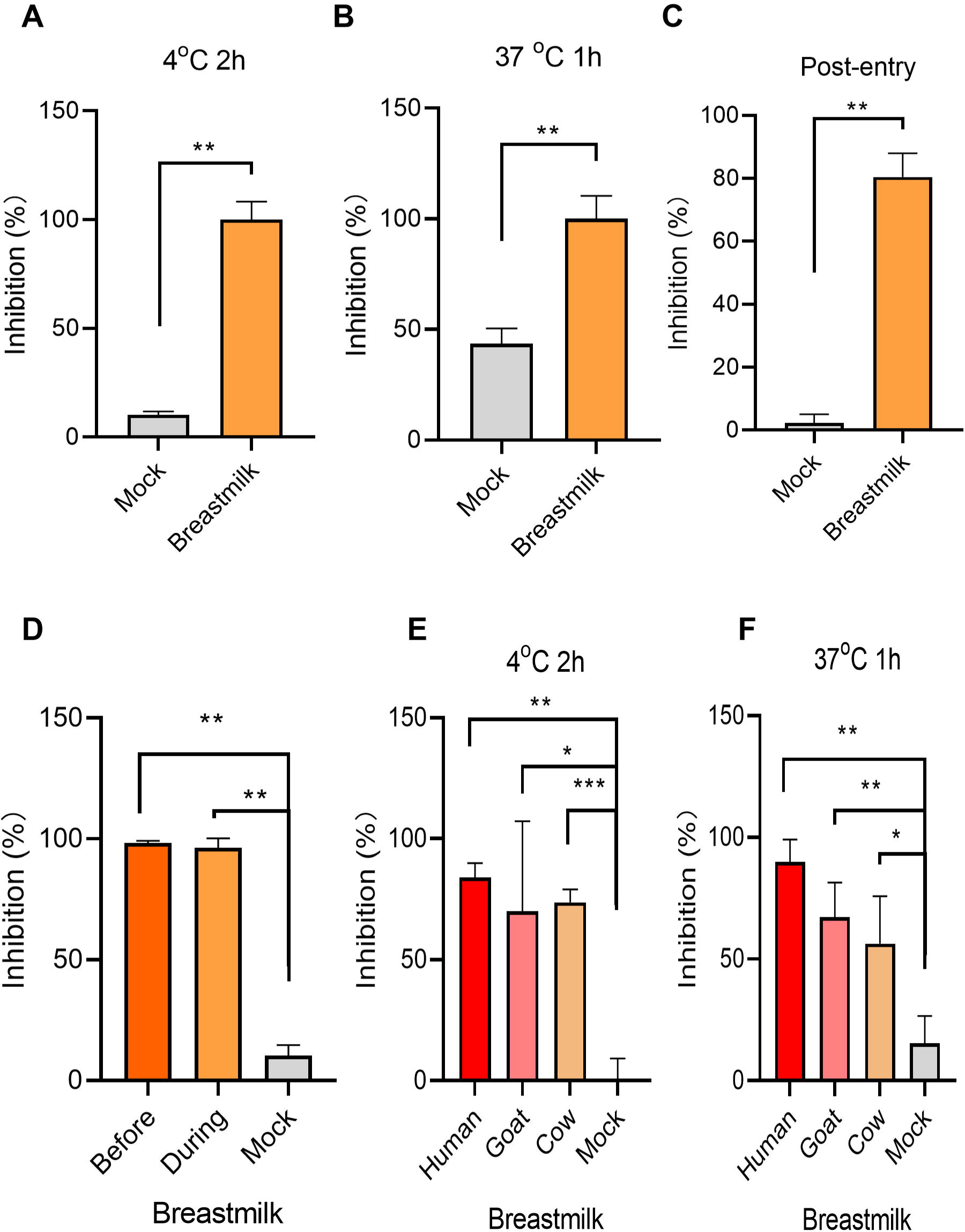
Human breastmilk inhibited viral attachment, entry and post-entry replication. Breastmilk at final concentration of 4 mg/ml was mixed with GX_P2V for 1h at room temperature. Vero E6 cells were incubated with the mixture at 4°C for 2h (MOI=10) (**A**) and 37°C for 1h (MOI=10) (**B**), respectively. After washing out of the free viruses, the cells were harvested and detected by RT-qPCR and normalized by GAPDH level. (**C**) Vero E6 cells were infected with GX_P2V (MOI=0.01) at 37°C for 1h. After washing out the free viruses, the cells were treated with breastmilk at final concentration of 4mg/ml for 72h. Viral RNA were quantified by RT-qPCR to test the post-entry replication. To test whether breastmilk binding to cell surface to block SARS-CoV-2 pseudovirus infection, breastmilk at final concentration of 4mg/ml was incubated with Vero E6 cells at 37°C for 1h. After washing out of the free breastmilk, the cells were infected with SARS-CoV-2 pseudovirus (650 TCID_50_/well) and cultured at 37°C for 24h (**D**). To test whether breastmilk inhibit viral attachment and entry, Vero E6 cells were incubated with the mixture of breastmilk and SARS-CoV-2 pseudovirus (650 TCID_50_/well) at 4°C for 2h (**E**) and 37°C for 1h (**F**), respectively. After washing out of the free viruses, Vero E6 cells were cultured at 37°C for 24h. SARS-CoV-2 pseudovirus infection was measured by luciferase assay. Values are shown as mean of triplicates ± SD, \**p*<0.05, \*\**p*<0.01, \*\*\**p*<0.001 by unpaired two-tailed t test.

## Discussion

In the present study, we demonstrated for the first time that whey protein from human breastmilk significantly inhibited the infection of SARS-CoV-2 and its related pangolin coronavirus (GX_P2V) in cells model study. Breastmilk could not only block viral attachment and entry, but also inhibit post-entry viral replication.

Because of biosafety consideration, we did not directly use wild live SARS-CoV-2, but used pseudovirus and GX_P2V in the present study. Studies demonstrate that pseudovirus is suitable for neutralization assay against SARS-CoV-2(14, 19). GX_P2V, which was identified from the pangolins recently, shared 92.2% amino acid of spike protein with SARS-CoV-2 and also infected the cells through binding the angiotensin-converting enzyme 2 (ACE2) receptor(13, 15). This virus would be a suitable alternative model for SARS-CoV-2 related research and studied at biosafety level-2, which could be studied in many labs. In our present study, we found that human breastmilk inhibited the infectivity of both SARS-CoV-2 pseudovirus and GX_P2V, with EC_50_<0.2mg/ml. The anti-coronavirus property of human breastmilk was confirmed in *in vitro* study with two different but closely related coronavirus and also two different cell lines of Vero E6 and A549 cells. Thus, it is reasonable to deduce that human breastmilk has the property of prevent SARS-CoV-2 infection and replication.

As we all known, human breastmilk contains different concentrations of the components among different species, such as cow and goat(11). They also have different antimicrobial activity(11). In the present study, we also revealed that whey protein from cow and goat inhibited the infectivity of SARS-CoV-2 pseudovirus and GX_P2V, although the inhibition efficiency was relatively lower compared to that of human whey protein. These results indicated that human whey protein has high concentration of antiviral factors than those from other species.

It was reported that human milk is rich in LF (3-5g/L in mature milk), which is 10-100 fold higher than that in cow and goat milk(20). In addition, LF is the most prominent antimicrobial component in milk(11). However, our results showed that either hLF or bLF showed limited anti-coronavirus activity compared the skimmed milk, indicating that LF is less likely to be a key component to inhibit SARS-CoV-2. Since milks of cow and goat contain much lower concentration (0.02–0.2mg/mL) of LF than human breastmilk (1-7mg/ml), the findings that milks of cow and goat inhibited the infectivity of SARS-CoV-2 also suggest that LF is not the active component to inactivate SARS-CoV-2(21). Besides LF, specific immunoglobulins are also important for the host defense system(18). It was reported that IgA antibody from the recovery patients blocks the interaction of SARS-CoV-2 and ACE2(18). As the human breastmilk samples were collected before the emergence of COVID-19, the breastmilk donors should have not been infected with SARS-CoV-2. Moreover, SARS-CoV-2 specific IgA was negative in the human breastmilk samples used in the present study and antibodies directed against human IgA did not offset the inhibition of human breastmilk on the infectivity of SARS-CoV-2 pseudovirus and GX_P2V. Therefore, the inhibition of whey protein on the infectivity of SARS-CoV-2 is not due to the IgA antibody. Together, these results suggested that other mechanism may be responsible for anti-SARS-CoV-2 activity of whey protein, which merits further study.

In conclusion, we uncovered that whey protein from human breastmilk inhibits SARS-CoV-2 virus infection in cultured cells. It is worth to identify the key factors for further antiviral drug development.

