## supplemental table 1 for "The effect of whey protein on viral infection and replication of SARS-CoV-2 and pangolin coronavirus in vitro"

**Supplementary tables**

**Supplementary Table 1:** **Primers used in the study.**

| **Primer name** | **Sequence** |
| --- | --- |
| CoV-F1 | GGTGATTGCCTTGGTGATATTG |
| CoV-R1 | GCAAGTAGTGCAGAAGTGTATTG |
| CoV-Probe | TCTGTGAGCAAAGGCGGTAGAACC（5-FAM，3-TAMRA） |
| GAPDH-q-F | AGCCTCAAGATCATCAGCAATG |
| GAPDH-q-R | ATGGACTGTGGTCATGAGTCCTT |
| GAPDH-q-probe | CCAACTGCTTAGCACCCCTGGCC（5-FAM，3-TRAMA） |
